# A heuristic method for fast and accurate phasing and imputation of single nucleotide polymorphism data in bi-parental plant populations

**DOI:** 10.1101/330027

**Authors:** Serap Gonen, Valentin Wimmer, R. Chris Gaynor, Ed Byrne, Gregor Gorjanc, John M. Hickey

**Affiliations:** The Roslin Institute and Royal (Dick) School of Veterinary Studies, University of Edinburgh, Easter Bush Research Centre, Midlothian EH25 9RG, UK; KWS SAAT SE, Grimsehlstr. 31, 37574 Einbeck, Germany; KWS-UK Ltd, 56 Church Street, Thriplow, Hertfordshire, SG8 7RE, UK

## Abstract

This paper presents a new heuristic method for phasing and imputation of genomic data in diploid plant species. Our method, called AlphaPlantImpute, explicitly leverages features of plant breeding programs to maximise the accuracy of imputation. The features are a small number of parents, which can be inbred and usually have high-density genomic data, and few recombinations separating parents and focal individuals genotyped at low-density (i.e. descendants that are the imputation targets). AlphaPlantImpute works roughly in three steps. First, it identifies informative low-density genotype markers in parents. Second, it tracks the inheritance of parental alleles and haplotypes to focal individuals at informative markers. Finally, it uses this low-density information as anchor points to impute focal individuals to high-density.

We tested the imputation accuracy of AlphaPlantImpute in simulated bi-parental populations across different scenarios. We also compared its accuracy to existing software called PlantImpute. In general, AlphaPlantImpute had better or equal imputation accuracy as PlantImpute. The computational time and memory requirements of AlphaPlantImpute were tiny compared to PlantImpute. For example, accuracy of imputation was 0.96 for a scenario where both parents were inbred and genotyped at 25,000 markers per chromosome and a focal F_2_ individual was genotyped with 50 markers per chromosome. The maximum memory requirement for this scenario was 0.08 GB and took 37 seconds to complete.

## Introduction

This paper presents a new heuristic method for phasing and imputation of single nucleotide polymorphism (SNP) array data in diploid plant species. High-density SNP array data in plant breeding populations is increasingly valuable for genomic selection and for identifying regions of the genome that underlie traits of interest in genome-wide association studies. The accuracy of genomic selection and power of association studies increases with the number of individuals and with the density of SNP markers. However, the cost of genotyping many individuals at high-density is high. This high cost is a barrier to the adoption of genomic selection in plant breeding programs where the number of selection candidates in each cycle can be very large. An effective strategy to overcome this cost barrier is to genotype a proportion of the population at high-density, phase their genotypes, and use this data for imputation of large numbers of individuals genotyped at low-density (Jacobson et al., 2014, 2015; Gorjanc et al., 2017a; b). This strategy has been widely adopted in livestock and human populations, partly because genotype imputation tools that work well in these populations are widely available (Kong et al., 2008; Howie et al., 2009; Druet and Georges, 2010; Li et al., 2010; Sargolzaei et al., 2011; Hickey et al., 2011; Cleveland and Hickey, 2013; Hickey and Kranis, 2013; VanRaden et al., 2015; O’Connell et al., 2016; Loh et al., 2016; Antolín et al., 2017).

Bi-parental populations that are widely used in plant breeding have four features that make them ideal for imputation. First, they are derived from only two parents. High-density genotyping of the two parents and low-density genotyping of focal individuals (i.e., descendants that are the imputation targets) is an effective low-cost strategy in these populations. Second, the number of meiosis separating parents and focal individuals is small. This means that parental haplotypes remain largely intact in focal individuals, which simplifies imputation. Third, they have well-known crossing structures that could be informative for imputation, although the process of selfing or the creation of doubled haploids can add complications that are not present in human and livestock settings. However, these “complications” can in certain situations empower imputation. Finally, parents that contribute to a bi-parental population are usually inbred. This means that they are homozygous at many loci and the majority of their genome is phased *de facto*.

A recent simulation study demonstrated that achieving high imputation accuracies could empower genomic selection in bi-parental populations (Gorjanc et al., 2017a; b). The high imputation accuracies with SNP array data were achieved using the PlantImpute software (Nettelblad et al., 2009; Hickey et al., 2015). The main drawback of PlantImpute is that it has large computational requirements in terms of time and memory. This makes it impractical for routine use in breeding programs. Existing software for imputation in livestock or human populations do not have large computational requirements. However, software for imputation in livestock or human populations are not designed to leverage features of plant breeding programs, and in some cases, cannot work where selfing and bi-sexuality is common. To our knowledge, existing imputation software for plant breeding programs (e.g., (Swarts et al., 2014)) are not explicitly designed for imputation of SNP array genotypes in bi-parental populations.

This paper presents a new heuristic method, called AlphaPlantImpute, for phasing and imputation of SNP array data in diploid plant species. AlphaPlantImpute works roughly in three steps. First, it identifies markers fully or partially informative for parent-of-origin. Second, it tracks the inheritance of parental alleles and haplotypes to focal individuals at informative markers. Finally, it uses this low-density information as anchor points to impute focal individuals to high-density.

We tested the accuracy of AlphaPlantImpute in simulated bi-parental populations across different scenarios. These scenarios varied in the levels of inbreeding in the parents, the number of selfing events separating parents and focal individuals, the chromosome size (i.e. recombination rate) and the number of markers on the low-density array. We calculated the accuracy of imputation within each scenario as the correlation between the true and imputed genotypes. In general, AlphaPlantImpute gave excellent accuracy of imputation and typically outperformed or performed equally as well as PlantImpute for the accuracy of imputation. The computational time and memory requirements of AlphaPlantImpute were always tiny compared to that of PlantImpute.

## Materials and methods

### Definitions

A focal individual is an individual that is to be imputed. A fully informative marker is one where the two parents have opposing homozygous genotypes, i.e., genotypes 0 and 2 (note that the method is agnostic of which allele is the reference allele). A partially informative marker is where one parent is homozygous and the other is heterozygous. Markers where parents are fixed for the same allele or where both parents are heterozygous are uninformative. The high-density (**HD**) array is the array at which parents have genotypes and is the target array for imputation. In our test datasets, the HD array consisted of 25,000 SNP markers. The low-density (**LD**) array is the array at which focal individuals have genotypes. We tested eight LD arrays (see below), all of which were nested subsets of the HD array.

### Description of the method

We present a new heuristic method, called AlphaPlantImpute, for phasing and imputation of SNP array data in diploid plant species. In detail, our method has five steps: (1) Identify markers that are informative for parent-of-origin of alleles in focal individuals; (2) Infer the most likely linked alleles at two markers; (3) Phase and assign parent-of-origin for focal individual’s alleles; (4) Impute focal individual to high-density using low-density anchors captured in step 3; and (5) Impute markers in recombined regions. Impute markers adjacent to recombination locations. Step 1 is the only step applied to groups of focal individuals together. Steps 2, 3, 4 and 5 are applied for each focal individual separately. A description of the definitions used and of each step is given below and a schematic is given in Figure 1 (a more detailed schematic is given in Supplementary Figure 1).

**Figure 1.**
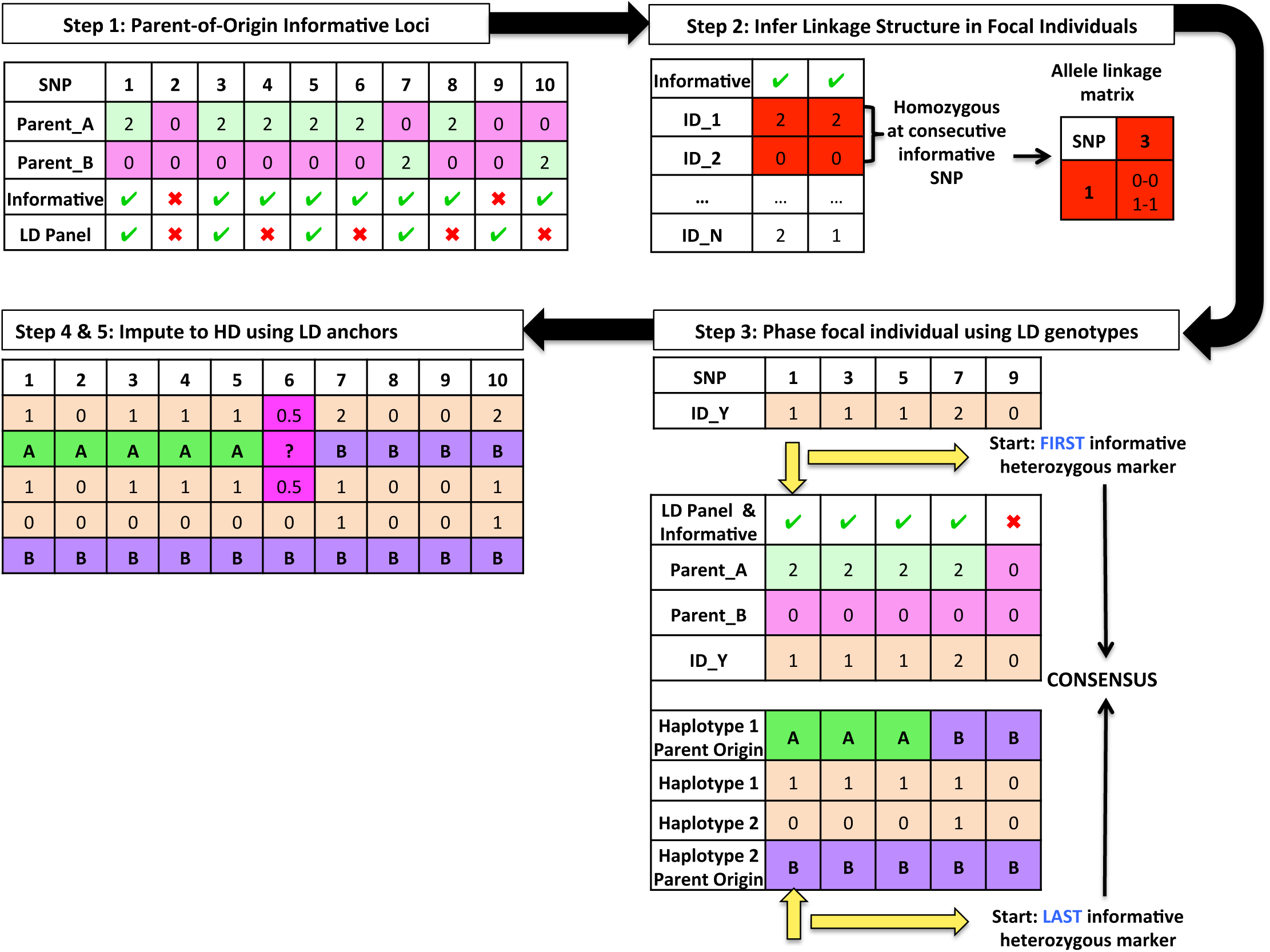
Schematic of heuristic algorithm of AlphaPlantImpute

### Method steps

#### Step 1: Identify informative low-density markers in parents

In the first step we determine which low-density markers are fully or partially informative in parents, which is used in the following steps to infer parent-of-origin of phased alleles in focal individuals. For example, in Figure 1 eight of the ten markers on the HD array genotyped in the parents are fully informative and two (markers 2 and 9) are uninformative. Of the ten HD markers, five (markers 1, 3, 5, 7, 9) are also on the LD array, which was used to genotype focal individuals. Of these five LD markers, four are informative and one (marker 9) is uninformative.

#### Step 2: Infer the most likely linked alleles at two markers

In the second step we infer the most likely linked alleles at two markers for all pairs of informative markers, which is used in the following steps to phase heterozygous markers in focal individuals. If parent haplotypes are inherited directly without recombination, the most likely linked alleles at two markers recover the parent haplotypes. When this is not the case, the most likely linked alleles at two markers indicate a potential recombination hotspot or marker map error for the population. For each pair of informative markers we perform three steps.

2a) First, identify focal individuals that are homozygous at the first and the second marker.

2b) Second, count the number of times focal individuals have genotype:

- 0 for the first and 0 for the second marker (diplotype 0-0),
- 0 for the first and 2 for the second marker (diplotype 0-2),
- 2 for the first and 0 for the second marker (diplotype 2-0), and
- 2 for the first and 2 for the second marker (diplotype 2-2).

2c) Third, compare the count of 0-0 to 0-2 and of 2-2 to 2-0. If the count of 0-0 is higher than 0-2 and 2-2 is higher than 2-0, then the 0 (1) allele at the first marker is commonly linked to the 0 (1) allele at the second marker. If the count of 0-2 is higher than 0-0 and 2-0 is higher than 2-2, then the 0 (1) allele at the first marker is commonly linked to the 1 (0) allele at the second marker. For example, in Figure 1 2-2 and 0-0 are the two most frequent diplotypes at markers 1 and 3, which suggests the most likely linked alleles are 1-1 and 0-0.

#### Step 3: Phase and assign parent-of-origin for focal individual’s alleles

In the third step we phase alleles in focal individuals and assign their parent-of-origin. We perform this first for the homozygous markers and then for the heterozygous markers.

##### 3a) Phase homozygous markers

We phase alleles at homozygous markers as the 0 allele for both haplotypes when the genotype is 0 and as the 1 allele when the genotype is 2. For example, in Figure 1 the focal individual ID_Y has genotype 2 for marker 7 and we phase it as the 1 allele for both haplotypes.

##### 3b) Assign parent-of-origin to alleles at homozygous markers

We assign parent-of-origin for phased alleles in the step 3a based on the informative markers in the step 1. For example, in Figure 1 marker 7 is informative. At this marker, the Parent_A has the 0 allele, while the Parent_B has the 1 allele. Focal individual ID_Y has genotype 2, which suggests that both of the 1 alleles were inherited from the Parent_B. Focal individual ID_Y is also homozygous at marker 9, with genotype 0, but this marker is not informative and we cannot assign parent-of-origin to phased alleles.

##### 3c) Phase heterozygous marker

We phase alleles at heterozygous markers iteratively based on the most likely linked alleles in the step 2. Specifically, we perform four steps. We start at the first heterozygous marker. For example, in Figure 1 the first marker for which the focal individual ID_Y is heterozygous is marker 1.

3c1) First, phase the first heterozygous marker randomly as the 1 allele for the first haplotype and the 0 allele for the second haplotype.

3c2) Second, phase the second heterozygous marker based on the the most likely linked alleles in the step 2. For example, in Figure 1 the second heterozygous marker is marker 3. Information from the most likely linked alleles suggest that the 0 (1) allele at marker 1 is linked to the 0 (1) allele at marker 3. Using this information, we phase marker 3 alleles of ID_Y as the 1 allele for the first haplotype and the 0 allele for the second haplotype. We continue moving from left-to-right until the last heterozygous marker is phased.

3c3) Third, we repeat steps 3c1 and 3c2, but this time starting from the last heterozygous marker and progressing to the first heterozygous marker.

3c4) Finally, we derive a consensus between the haplotypes derived from moving left-to-right and right-to-left along the chromosome. If they disagree, set the consensus haplotypes to missing. If only one is filled, set the consensus haplotype to the filled information.

##### 3d) Assign parent-of-origin to alleles at heterozygous marker

We assign parent-of-origin for phased alleles in the step 3c based on the informative markers in the step 1. For example, in Figure 1 focal individual ID_Y is heterozygous at marker 1. At this marker, the 1 allele on ID_Y’s first haplotype is inherited from Parent_A and the 0 allele on ID_Y’s second haplotype is inherited from Parent_B. If the marker is partially informative, we assign both the parent-of-origin and the haplotype-of-origin (i.e., first or second haplotype of the parent that is heterozygous for that marker).

#### Step 4: Impute focal individual to high-density using anchors from the step 3

##### 4a) Fill uninformative homozygous markers

For uninformative homozygous markers at HD that are not genotyped in the focal individual at LD, we phase and impute the focal individual with the parental information. For example, in Figure 1 both parents have genotype 0 for marker 2, so focal individual ID_Y is imputed as genotype 0.

##### 4b) Assign parent-of-origin to HD marker alleles

For markers on the HD array, assign parent-of-origin to marker alleles based on the parent-of-origin assignment of the two nearest marker alleles on the LD array. For example, in Figure 1 marker 6 is not genotyped on the LD array but the two neighbouring markers 5 and 7 are genotyped on the LD array. We have assigned the second haplotype of focal individual ID_Y to Parent_B for both markers 5 and 7. We therefore also assign marker 6 to Parent_B for the second haplotype. We have assigned the first haplotype of focal individual ID_Y to Parent_A for marker 5 and to Parent_B for marker 7. We conclude that there was a potential recombination around marker 6 at the first haplotype and we do not assign parent-of-origin for this allele.

##### 4c) Phase and impute HD markers using parent-of-origin assignment from step 4b

For HD markers with assigned parent-of-origin in step 4b, we phase the allele inherited from that parent for the haplotype of the focal individual. If we have phased both alleles at a marker, we impute the genotype as the sum of the two alleles on the two haplotypes of the focal individual. If parent-of-origin has not been assigned for one or both alleles of the focal individual, we leave the genotype as missing.

#### Step 5. Impute markers in recombined regions

We phase and impute missing HD markers in potentially recombined regions in one of two ways. We either (1) impute expected genotype dosage as the average of the alleles of the two parents; or (2) phase and impute using information from a genetic or physical map. For (2), we first identify the two closest neighbouring markers that were informative and phased, second use the distance between these two markers as a weight to phase the missing alleles as the weighted average of the alleles of the two parent haplotypes, and third impute expected genotype dosage as in (1).

### Implementation

We have implemented the method in a program called AlphaPlantImpute, which is controlled by a specification file that contains some user specified thresholds and the addresses of input files. The required input data are membership of individuals to the bi-parental populations, HD genotypes for parents, and LD genotypes of focal individuals. The output data are imputed genotypes, phased haplotypes, inferred parent-of-origin for focal individual haplotypes, and information on whether a marker is informative. AlphaPlantImpute implements some data editing checks, which are described in the user manual.

### Examples of implementation: Description of datasets

To test the imputation accuracy of AlphaPlantImpute, testing datasets of a subset of the scenarios described in Hickey et. al. 2015 were simulated. This enabled the comparison of AlphaPlantImpute with PlantImpute without re-running PlantImpute with its large computational cost. Although the simulation design is largely a replication of that in Hickey et. al. 2015, a brief description of the general structure and simulation method of the different scenarios tested is given below for completeness.

### Simulation of genomic data

Sequence data for 100 base haplotypes for a single chromosome were simulated using the Markovian Coalescent Simulator (Chen et al., 2009) and AlphaSim (Faux et al., 2016). The base haplotypes were 10^8^ base pairs in length, with a per site mutation rate of 1.0×10^−8^ and a per site recombination rate that varied across scenarios. The different recombination rates simulated were 0.25×10^−8^, 0.5×10^−8^, 1.0×10^−8^, 1.5×10^−8^, 2.0×10^−8^, 3.0×10^−8^, and 4.0×10^−8^, resulting in chromosome sizes of 25, 50, 100, 150, 200, 300, and 400 centiMorgans (cM), respectively. The effective population size (N_e_) was set at specific points during the simulation to mimic changes in N_e_ in a crop such as maize *(Zea mays L.)*. These set points were: 100 in the base generation, 1000 at 100 generations ago, and 10,000 at 2000 generations ago, with linear changes in between. The resulting whole-chromosome haplotypes had approximately 80,000 segregating sites in total.

### Simulation of a pedigree

A pedigree of 11,266 individuals was constructed. The pedigree was initiated from six outbred founders (A, B, C, D, E, F). These six founders were crossed to generate the founder bi-parental populations (AxB, CxD, ExF). These founder bi-parental populations were selfed to F_1_, F_2_, F_4_, F_10_, or F_20_, resulting in different levels of inbreeding in the parents. To properly propagate the residual heterozygosity in these parents, they were crossed to generate 100 pairs of F_1_ individuals. F_1_ individuals were selfed to generate 100 F_2_ individuals. F_2_ individuals were selfed to generate 100 F_3_ individuals, and selfing continued through to F_10_. The focal individuals (i.e. descendants that were the imputation targets) were F_2_, F_4_, F_6_, or F_10_ descendants.

In the base generation, individuals had their chromosomes sampled from the 100 base haplotypes. In subsequent generations the chromosomes of each individual was sampled from parental chromosomes with recombination. The recombination rate varied depending on the scenario resulting in chromosome sizes of 25, 50, 100, 150, 200, 300, and 400 centiMorgans (cM). Recombinations occurred with a 1% probability per cM and were uniformly distributed along the chromosome.

### Simulated SNP marker arrays

A single HD array of 25,000 SNP markers for the single chromosome was simulated. To test the effect of the number of markers on the LD array, eight LD arrays of 3, 5, 10, 20, 50, 100, 200, and 400 markers for the single chromosome were simulated. Arrays were constructed by aiming to select a set of markers that segregated in the parents and that were evenly distributed across the chromosome. All LD arrays were nested within each other and within the HD array.

### Scenarios

The imputation accuracy of AlphaPlantImpute and PlantImpute were compared in four different scenarios (scenario 1, 2, 3, and 4). Scenarios 1, 2, and 3 were the same as scenarios 2, 4, and 5 in Hickey et al. 2015. A description of all four scenarios is provided below. In all scenarios, focal individuals genotyped at LD were imputed to the single HD array of 25,000 SNP markers. Ten replications of each scenario were performed and the average of each replication is reported in the results.

Scenario 1: The effect of the number of selfing events separating parents and focal individuals. Parents were almost fully inbred (F_20_) and chromosomes were 100 cM in length. The accuracy of imputation was assessed for F_2_, F_4_, F_6_, and F_10_ focal individuals.

Scenario 2: The effect of the level of inbreeding in parents. Parents were F_1_, F_2_, F_4_, F_10_, or F_20_ and chromosomes were 100 cM in length. The accuracy of imputation was assessed for F_2_ focal individuals.

Scenario 3: The effect of chromosome size. Parents were fully inbred (F_20_) and the accuracy of imputation was assessed for F_2_ focal individuals. Chromosomes were 25, 50, 100, 150, 200, 300, or 400 cM in size.

Scenario 4: The effect of number of focal individuals in the bi-parental population. Parents were fully inbred (F_20_) and the accuracy of imputation was assessed for F_2_ focal individuals. Subsets of focal individuals were randomly selected from the 100 focal individuals to generate bi-parental population sizes of 1, 5, 10, 25, and 50 focal individuals.

### Analysis

Imputation was performed within each bi-parental population. Parents were assumed genotyped at HD and focal individuals were assumed genotyped at LD. The imputation accuracy was calculated for each focal individual as the correlation between the true and imputed genotypes. The precision in imputation accuracy was calculated as the log of the inverse of the variance in imputation accuracy within each bi-parental population.

## Results

For each scenario, we first present the imputation accuracy of AlphaPlantImpute and then compare it to PlantImpute (Nettelblad et al., 2009; Hickey et al., 2015).

### Effect of the number of markers on the low-density array

Increasing the number of LD markers increases the imputation accuracy of AlphaPlantImpute. Figure 2 plots the number of LD markers against the accuracy of imputation for F_2_ focal individuals of an F_20_ x F_20_ bi-parental cross. Figure 2 shows that increasing the number of LD markers from 3 to 20 SNP increased the average imputation accuracy from 0.85 to 0.96. Increasing the number of markers beyond 20 achieved only a slight increase in the accuracy of imputation from 0.96 with 20 markers to >0.99 with 400 markers.

**Figure 2.**
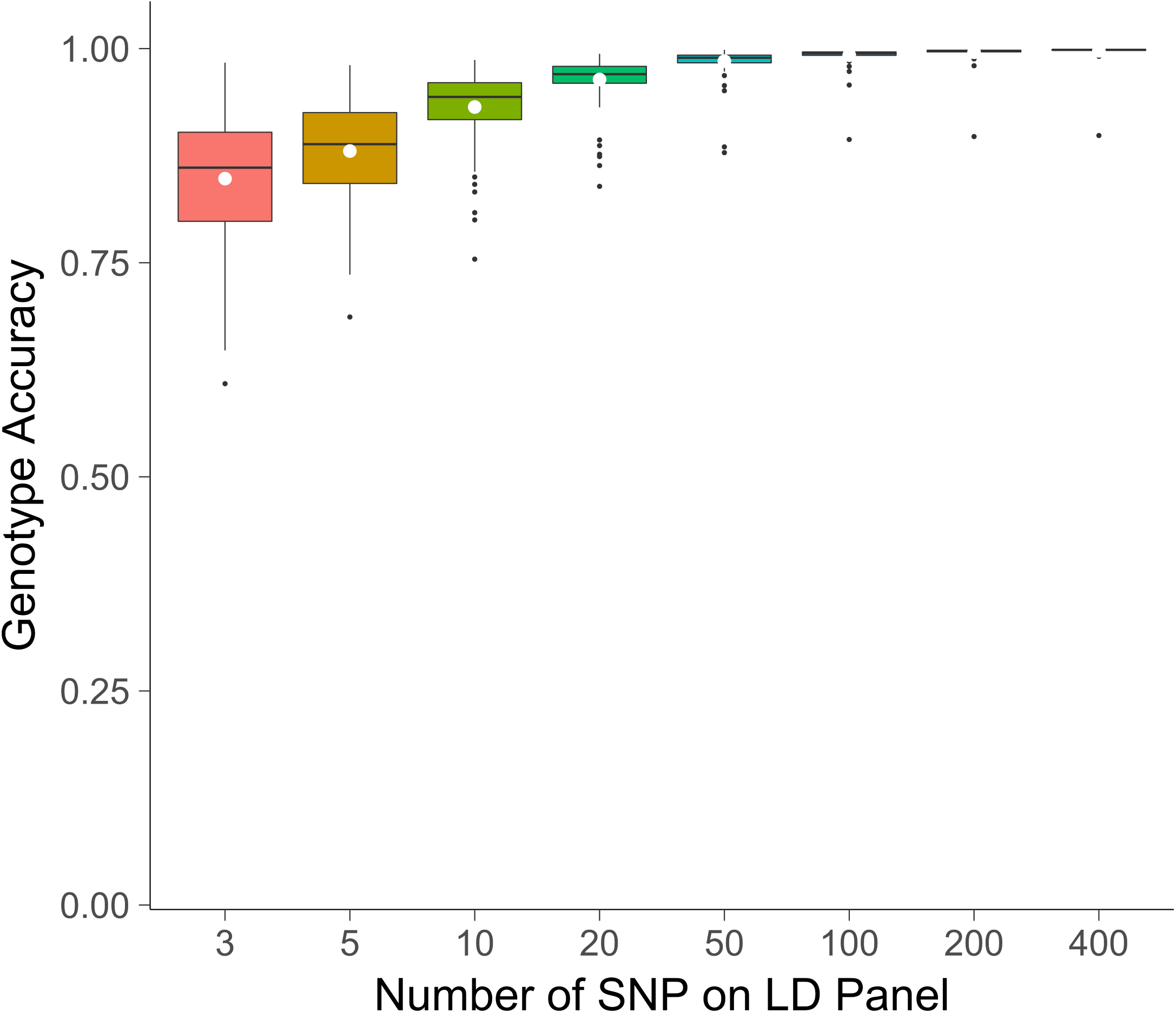
Effect of the number of SNP on the low-density panel. The number of SNP on the LD panel against the genotype imputation accuracy using AlphaPlantImpute for F_2_ focal individuals of a bi-parental cross where the parents are F_20_ inbred individuals.(c)

### Scenario 1: Effect of the number of selfing events separating parents and focal individuals

Increasing the number of selfing events separating parents and focal individuals slightly decreases the imputation accuracy of AlphaPlantImpute. Figure 3a plots the accuracy of imputation in F_2_, F_4_, F_6_ and F_10_ focal individuals of a bi-parental population where the parents were F_20_. Figure 3a shows that with 3 LD markers, the average imputation accuracy decreased from 0.85 for F_2_ focal individuals to 0.77 for F_10_ focal individuals. Increasing the number of LD markers beyond 10 markers mitigates the decrease in the average imputation accuracy between F_2_ focal individuals and F_10_ focal individuals. Figure 3a shows that with 20 LD markers, the average imputation accuracy decreased from 0.96 for F_2_ focal individuals to 0.95 for F_10_ focal individuals.

**Figure 3.**
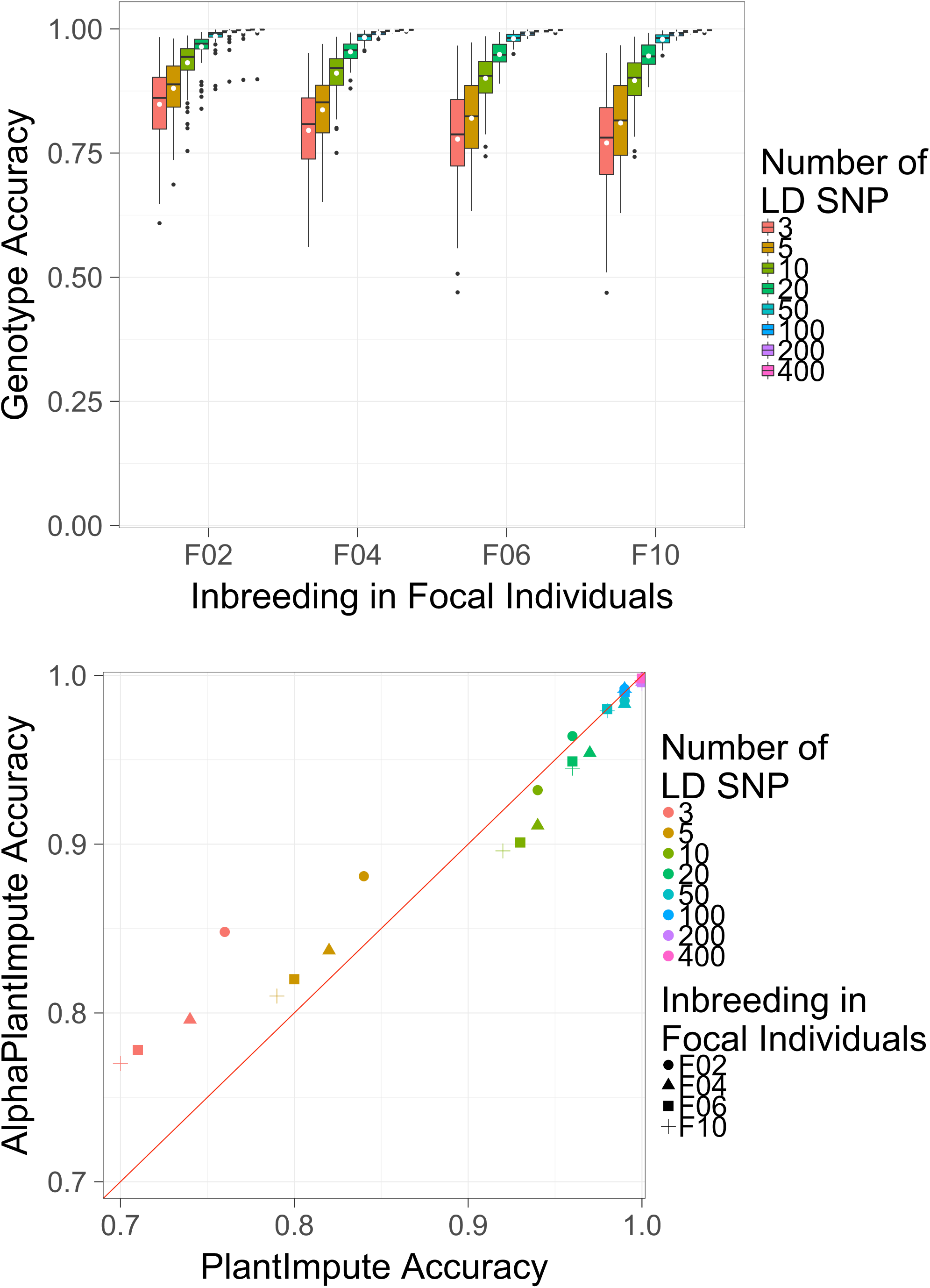
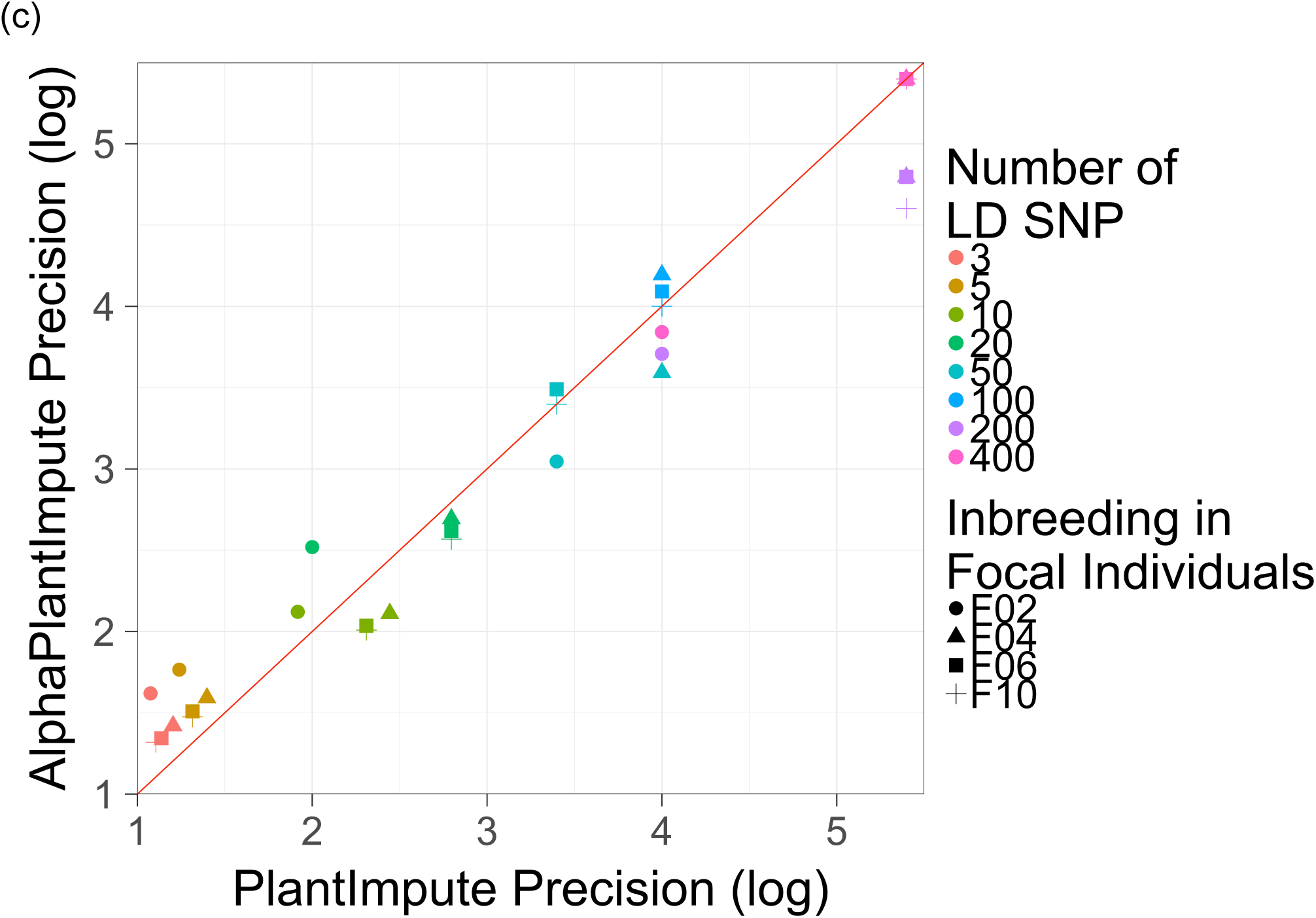
Effect of the level of inbreeding in focal individuals. (a)The genotype imputation accuracy using AlphaPlantImpute in F_2_, F_4_, F_6_ and F_10_ focal individuals from a bi-parental cross where the parents are F_20_ inbred individuals. (b)Comparison of the average genotype imputation accuracy using AlphaPlantImpute (y-axis) vs. PlantImpute (x-axis). The colours represent the different LD panels. The shapes represent the level of inbreeding in the focal individuals. The red diagonal line indicates when the accuracy of PlantImpute equals AlphaPlantImpute. Points above the line are when imputation accuracy is higher with AlphaPlantImpute and points below the line are when imputation accuracy is higher with PlantImpute. (c)Comparison of the precision in imputation accuracy using AlphaPlantImpute (y-axis) vs. using PlantImpute (x-axis). The colours represent the different LD panels. The shapes represent the level of inbreeding in the focal individuals. The red diagonal line indicates when the precision of PlantImpute equals AlphaPlantImpute. Points above the line indicate when the precision in accuracies is higher in AlphaPlantImpute and points below the line are when the precision in accuracies is higher in PlantImpute.

Regardless of the number of selfing events separating parents and focal individuals, the accuracy of imputation for AlphaPlantImpute was higher than for PlantImpute when the number of LD markers was low. Figure 3b plots the average imputation accuracy of AlphaPlantImpute on the y-axis and for PlantImpute on the x-axis. The colours represent the different number of LD markers and the shapes represent the number of selfing events separating the parents and the focal individuals. The red diagonal line indicates when the imputation accuracy of the two methods is equal. Points above the line indicate when the accuracy of imputation was higher for AlphaPlantImpute than for PlantImpute and visa versa. Figure 3b shows that with 3 LD markers, the average accuracy of imputation was 0.85 for AlphaPlantImpute and 0.76 for PlantImpute for F_2_ focal individuals and was 0.77 for AlphaPlantImpute and 0.70 for PlantImpute for F_10_ focal individuals.

For all numbers of selfing events separating parents and focal individuals, increasing the number of LD markers reduced and in some cases reversed the advantage of AlphaPlantImpute over PlantImpute. This was most obvious for F_10_ focal individuals for medium number of LD markers where the imputation accuracy with PlantImpute was slightly higher than with AlphaPlantImpute. Figure 3b shows that with 10 LD markers, the average imputation accuracy was 0.93 for AlphaPlantImpute and 0.94 for PlantImpute for F_2_ focal individuals and was 0.90 for AlphaPlantImpute and 0.92 for PlantImpute for F_10_ focal individuals. Increasing the number of LD markers beyond 100 markers meant that the average accuracy of imputation for AlphaPlantImpute equalled that for PlantImpute. Figure 3b shows that with 100 LD markers, the average imputation accuracy was 0.99 for both AlphaPlantImpute and PlantImpute for F_2_ focal individuals and for F_10_ focal individuals.

For all numbers of selfing events separating parents and focal individuals, the precision of imputation accuracy (i.e., consistency across focal individuals) for AlphaPlantImpute was higher than for PlantImpute when the number of LD markers was low. Figure 3c is similar to Figure 3b and plots the log of the precision of imputation accuracy for AlphaPlantImpute on the y-axis and PlantImpute on the x-axis. Points above the line indicate better precision (i.e. less variance) for AlphaPlantImpute than for PlantImpute, and vice versa. Figure 3c shows that with 3 LD markers, the precision of imputation was 1.62 for AlphaPlantImpute and 1.08 for PlantImpute for F_2_ focal individuals and was 1.32 for AlphaPlantImpute and 1.11 for PlantImpute for F_10_ focal individuals.

Figure 3c also shows that for medium number of LD markers, the precision of imputation accuracy for AlphaPlantImpute was higher than for PlantImpute for F_2_ focal individuals but was lower when the number of selfing events was higher. With 20 LD markers, the precision of imputation accuracy was 2.48 for AlphaPlantImpute and 2.00 for PlantImpute for F_2_ focal individuals and was 2.57 for AlphaPlantImpute and 2.80 for PlantImpute for F_10_ focal individuals. With the highest number of LD markers (400), the precision of imputation accuracy was 3.84 for AlphaPlantImpute and 4.00 for PlantImpute for F_2_ focal individuals and was 5.40 for both AlphaPlantImpute and PlantImpute for F_10_ focal individuals.

### Scenario 2: Effect of the level of inbreeding in parents

Increasing the level of inbreeding in the parents increases the imputation accuracy for AlphaPlantImpute. Figure 4a plots the accuracy of imputation in F_2_ focal individuals of a bi-parental population where the parents were F_1_, F_2_, F_4_, F_10_ or F_20_. Figure 4a shows that with 20 LD markers, the average imputation accuracy increased from 0.81 for F_1_ parents to 0.96 for F_20_ parents. Figure 4a also shows that increasing the level of inbreeding in the parents beyond F_4_ did not increase the average accuracy of imputation for F_2_ focal individuals. The average imputation accuracy with 20 LD markers was approximately 0.96 for F_2_ focal individuals when parents were F_4_, F_10_, and F_20_.

**Figure 4.**
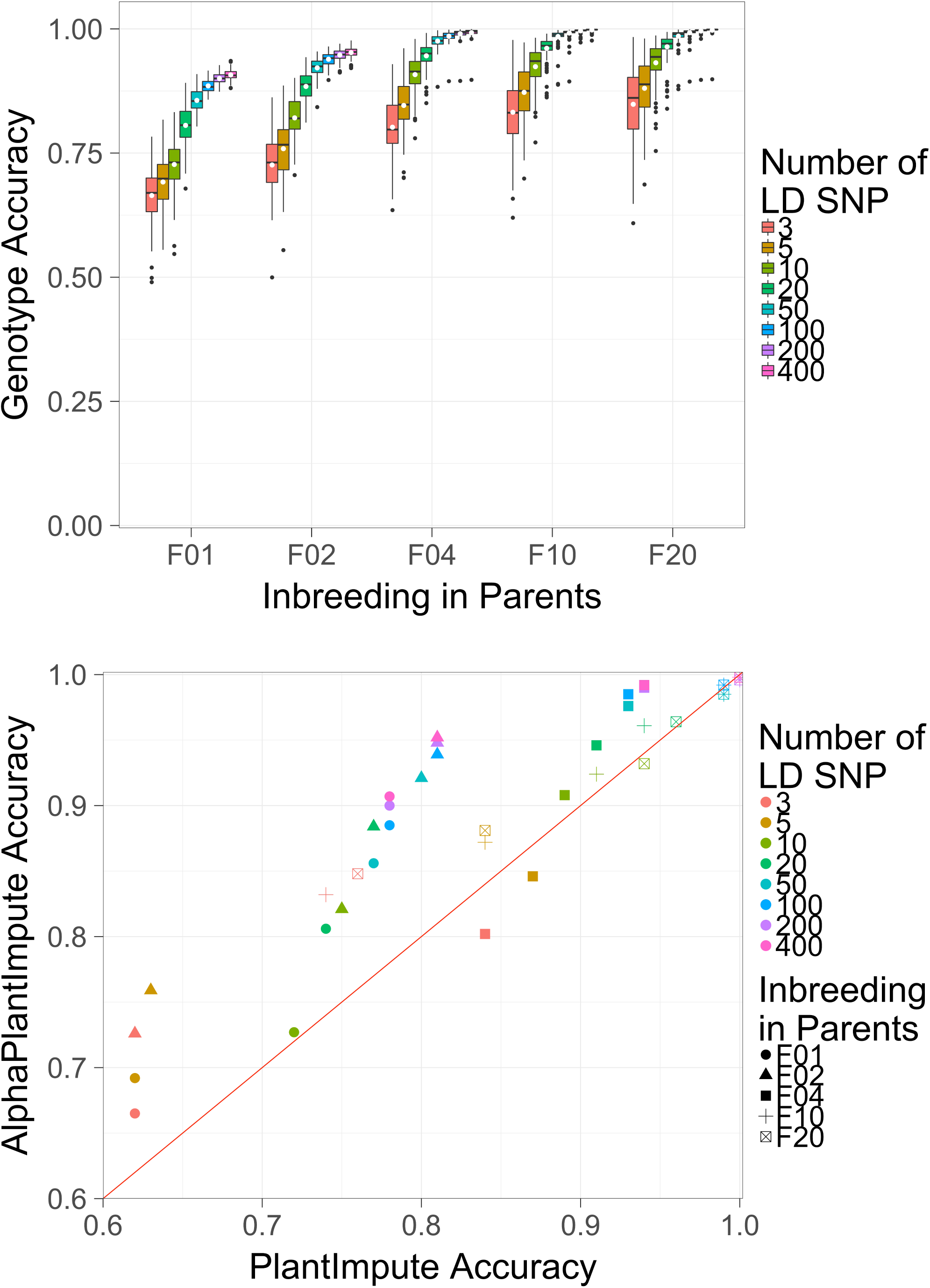
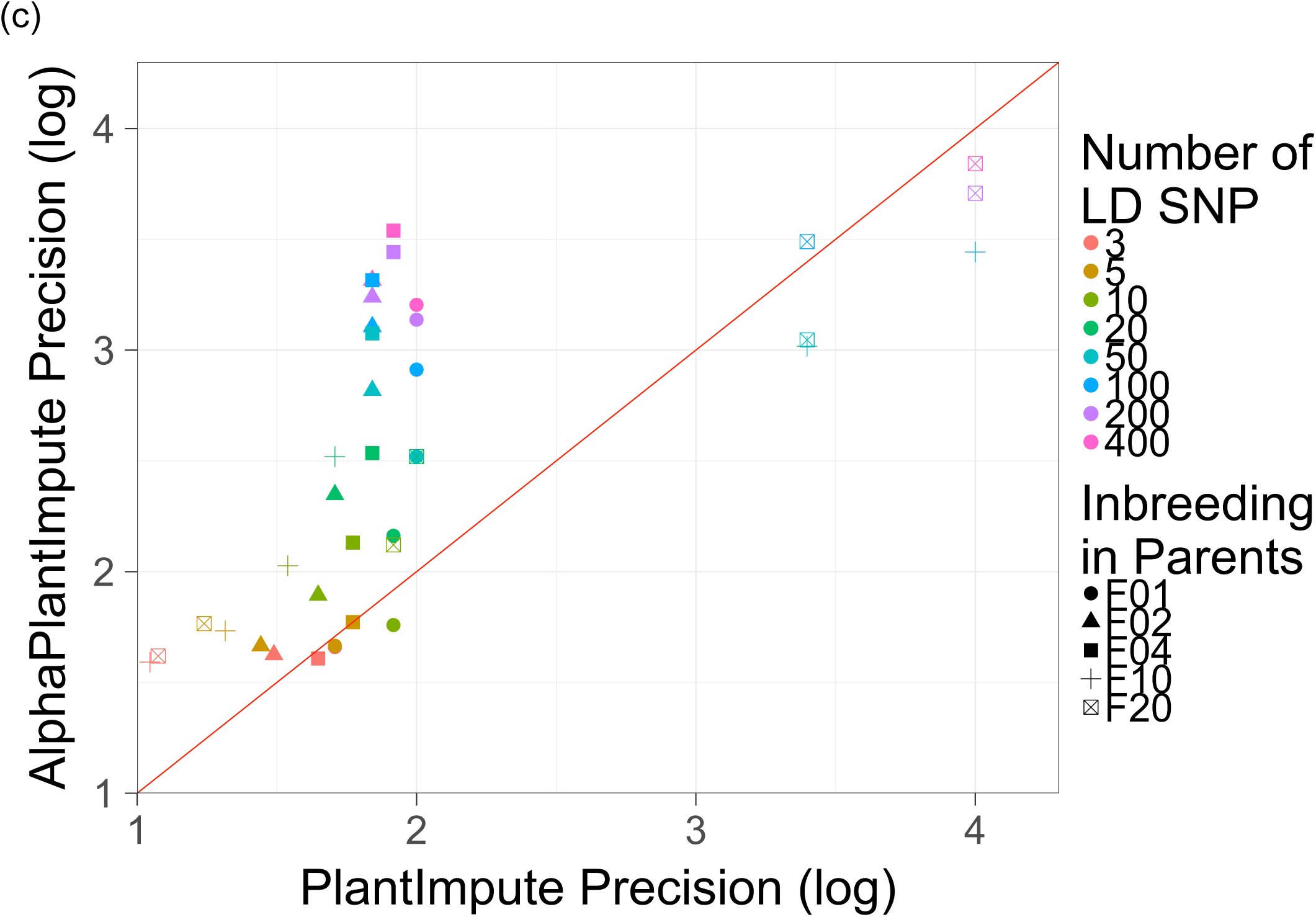
Effect of the level of inbreeding in parents. (a)The genotype imputation accuracy using AlphaPlantImpute in F_2_ focal individuals of a bi-parental cross where the parents are F_1_, F_2_, F_4_, F_10_ or F_20_. (b)Comparison of the average genotype imputation accuracy using AlphaPlantImpute (y-axis) vs. using PlantImpute (x-axis). The colours represent the different LD panels. The shapes represent the level of inbreeding in the parents. The red diagonal line indicates when the accuracy of PlantImpute equals AlphaPlantImpute. Points above the line are when imputation accuracy is higher with AlphaPlantImpute and points below the line are when imputation accuracy is higher with PlantImpute. (c)Comparison of the precision in imputation accuracy using AlphaPlantImpute (y-axis) vs. using PlantImpute (x-axis). The colours represent the different LD panels. The shapes represent the level of inbreeding in the parents. The red diagonal line indicates when the precision of PlantImpute equals AlphaPlantImpute. Points above the line indicate when the precision in accuracies is higher in AlphaPlantImpute and points below the line are when the precision in accuracies is higher in PlantImpute.

For all levels of inbreeding in the parents and all numbers of LD markers, the average imputation accuracy with AlphaPlantImpute was almost always higher than with PlantImpute. Figure 4b is similar to Figure 3b and plots the average imputation accuracy for AlphaPlantImpute on the y-axis and for PlantImpute on the x-axis. The shapes represent the level of inbreeding in the parents. Figure 4b shows that with 20 SNP LD markers, the average imputation accuracy was 0.81 for AlphaPlantImpute and 0.74 for PlantImpute for F_2_ focal individuals when parents were F_1_, 0.95 for AlphaPlantImpute and 0.91 for PlantImpute when parents were F_4_, and 0.96 for AlphaPlantImpute and 0.94 for PlantImpute when parents were F_10_. In two cases, the average imputation accuracy with PlantImpute was slightly higher than with AlphaPlantImpute. This was when parents were F_4_ and with 3 and 5 LD markers. The average imputation accuracy was 0.84 for AlphaPlantImpute and 0.80 for PlantImpute with 3 LD markers and was 0.87 for AlphaPlantImpute and 0.85 for PlantImpute with 5 LD markers.

For all levels of inbreeding in the parents and all numbers of LD markers, the precision of imputation accuracy with AlphaPlantImpute was almost always higher than with PlantImpute. Figure 4c is similar to 3c and plots the log of the precision of imputation accuracy for AlphaPlantImpute on the y-axis and PlantImpute on the x-axis. Figure 4c shows that with 20 LD markers, the precision of imputation accuracy was 2.16 for AlphaPlantImpute and 1.92 for PlantImpute for F_2_ focal individuals when parents were F_1_, 2.54 for AlphaPlantImpute and 1.84 for PlantImpute when parents were F_4_, and 2.52 for AlphaPlantImpute and 1.71 for PlantImpute when parents were F_10_. In a few cases, the precision of imputation accuracy for PlantImpute was slightly higher than AlphaPlantImpute. This was mainly when parents were F_20_ and with 50, 200, and 400 LD markers. The precision of imputation accuracy was 3.04 for AlphaPlantImpute and 3.40 for PlantImpute with 50 LD markers, was 3.71 for AlphaPlantImpute and 4.00 for PlantImpute with 200 LD markers, and was 3.84 for AlphaPlantImpute and 4.00 for PlantImpute with 400 LD markers.

### Scenario 3: Effect of chromosome size

Increasing the chromosome size (in cM) decreased the imputation accuracy for AlphaPlantImpute. This was most apparent when the number of LD markers was 10 or less. Figure 5a plots the imputation accuracy for seven chromosome sizes of 25, 50, 100, 150, 200, 300, and 400 cM for F_2_ focal individuals of a bi-parental population where the parents were F_20_. Figure 5a shows that with 3 LD markers, quadrupling the chromosome size from 25 cM to 100 cM decreased the average imputation accuracy from 0.95 to 0.85, and quadrupling from 100 cM to 400 cM decreased the average imputation accuracy from 0.85 to 0.55. The reduction in the imputation accuracy was less or non-existent when the number of LD markers was higher than 10. Figure 5a shows that the imputation accuracy was approximately 0.98 for all chromosome sizes when the number of LD markers was 50.

**Figure 5.**
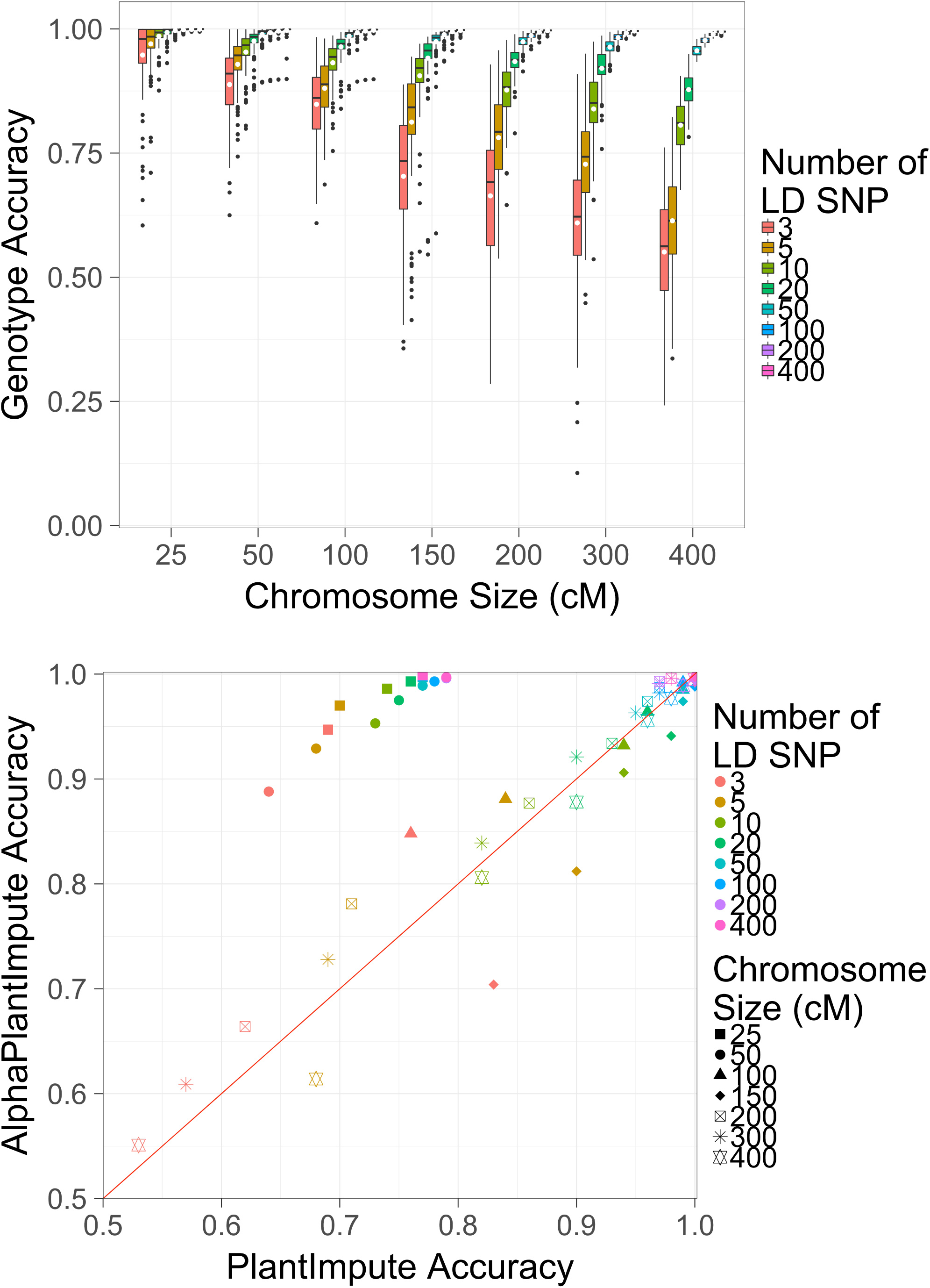
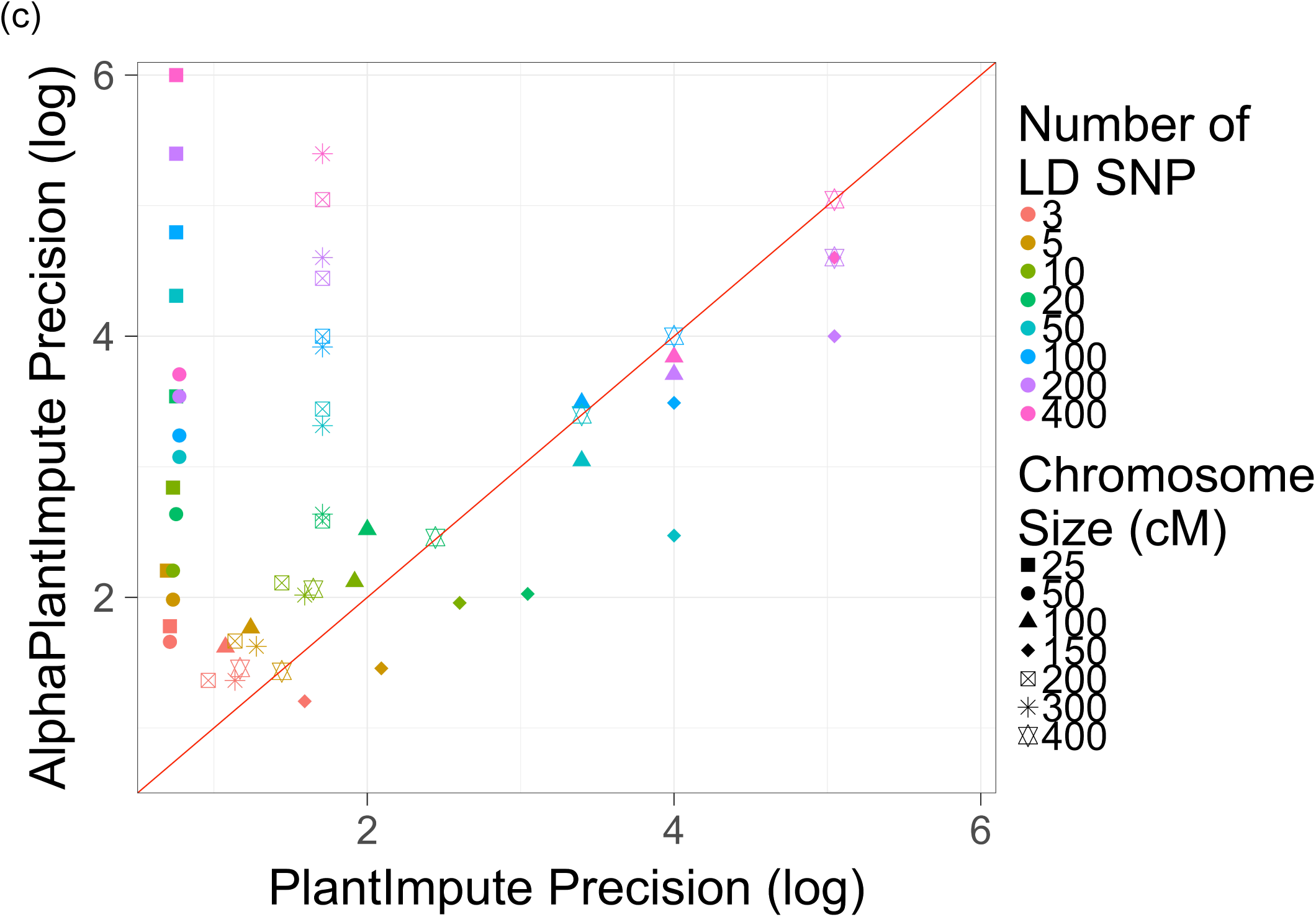
Effect of chromosome size. (a)The genotype imputation accuracy using AlphaPlantImpute in F_2_ focal individuals from a bi-parental cross of F_20_ parents against seven chromosome sizes of 25, 50, 100, 150, 200, 300, and 400 cM. (b)Comparison of the average genotype imputation accuracy using AlphaPlantImpute (y-axis) vs. using PlantImpute (x-axis). The colours represent the different LD panels. The shapes represent the chromosome size. The red diagonal line indicates when the accuracy of PlantImpute equals AlphaPlantImpute. Points above the line are when imputation accuracy is higher with AlphaPlantImpute and points below the line are when imputation accuracy is higher with PlantImpute. (c)Comparison of the precision in imputation accuracy using AlphaPlantImpute (y-axis) vs. using PlantImpute (x-axis). The colours represent the different LD panels. The shapes represent the chromosome size. The red diagonal line indicates when precision of PlantImpute equals AlphaPlantImpute. Points above the line indicate when the precision in accuracies is higher in AlphaPlantImpute and points below the line are when the precision in accuracies is higher in PlantImpute.

When the chromosome size was 300 cM or less, the average imputation accuracy was higher for AlphaPlantImpute than for PlantImpute. Figure 5b is similar to Figure 3b and plots the average imputation accuracy for AlphaPlantImpute on the y-axis and for PlantImpute on the x-axis. The shapes represent the chromosome sizes. Figure 5b shows that with 3 LD markers, the average imputation accuracy was 0.95 for AlphaPlantImpute and 0.69 for PlantImpute when the chromosome size was 25 cM and was 0.61 for AlphaPlantImpute and 0.57 for PlantImpute when the chromosome size was 300 cM. The exception to this was when the chromosome size was 150 cM, where the average imputation accuracy was 0.70 for AlphaPlantImpute and 0.83 for PlantImpute. When the chromosome size was 400 cM the average imputation accuracy was 0.55 for AlphaPlantImpute and 0.51 for PlantImpute when 3 LD markers were used but was 0.61 for AlphaPlantImpute and 0.68 for PlantImpute when 5 LD markers were used.

For all chromosome sizes and numbers of LD markers, the precision of imputation accuracy for AlphaPlantImpute was generally higher than for PlantImpute. Figure 5c is similar to Figure 3c and plots the precision of imputation accuracy for AlphaPlantImpute on the y-axis and for PlantImpute on the x-axis. Figure 5c shows that with 3 LD markers, the precision of imputation accuracy was 0.71 for AlphaPlantImpute and 1.78 for PlantImpute when the chromosome size was 25 cM, was 1.08 for AlphaPlantImpute and 1.62 for PlantImpute when the chromosome size was 100 cM and was 1.59 for AlphaPlantImpute and 1.20 for PlantImpute when the chromosome size was 400 cM. The exception to this was when the chromosome size was 150 cM, where the precision of imputation accuracy was 1.17 for AlphaPlantImpute and 1.46 for PlantImpute.

### Scenario 4: Effect of the number of focal individuals in the bi-parental population

Increasing the number of focal individuals in the bi-parental population slightly increased the imputation accuracy for AlphaPlantImpute. This was most apparent when the number of LD markers was low. Figure 6 plots the accuracy of imputation for F_2_ focal individuals of an F_20_ x F_20_ bi-parental cross with 1, 5, 10, 25, 50 or 100 focal individuals. Figure 6 shows that increasing the number of focal individuals from 5 to 100 increased the average imputation accuracy from 0.83 to 0.85 when 3 LD markers were used. Figure 6 also shows that when the 10 or more LD markers were used, increasing the number of focal individuals had no effect on the imputation accuracy. When the number of LD markers was 400, the average imputation accuracy was 0.96 with 5 or 100 focal individuals in the bi-parental population.

**Figure 6.**
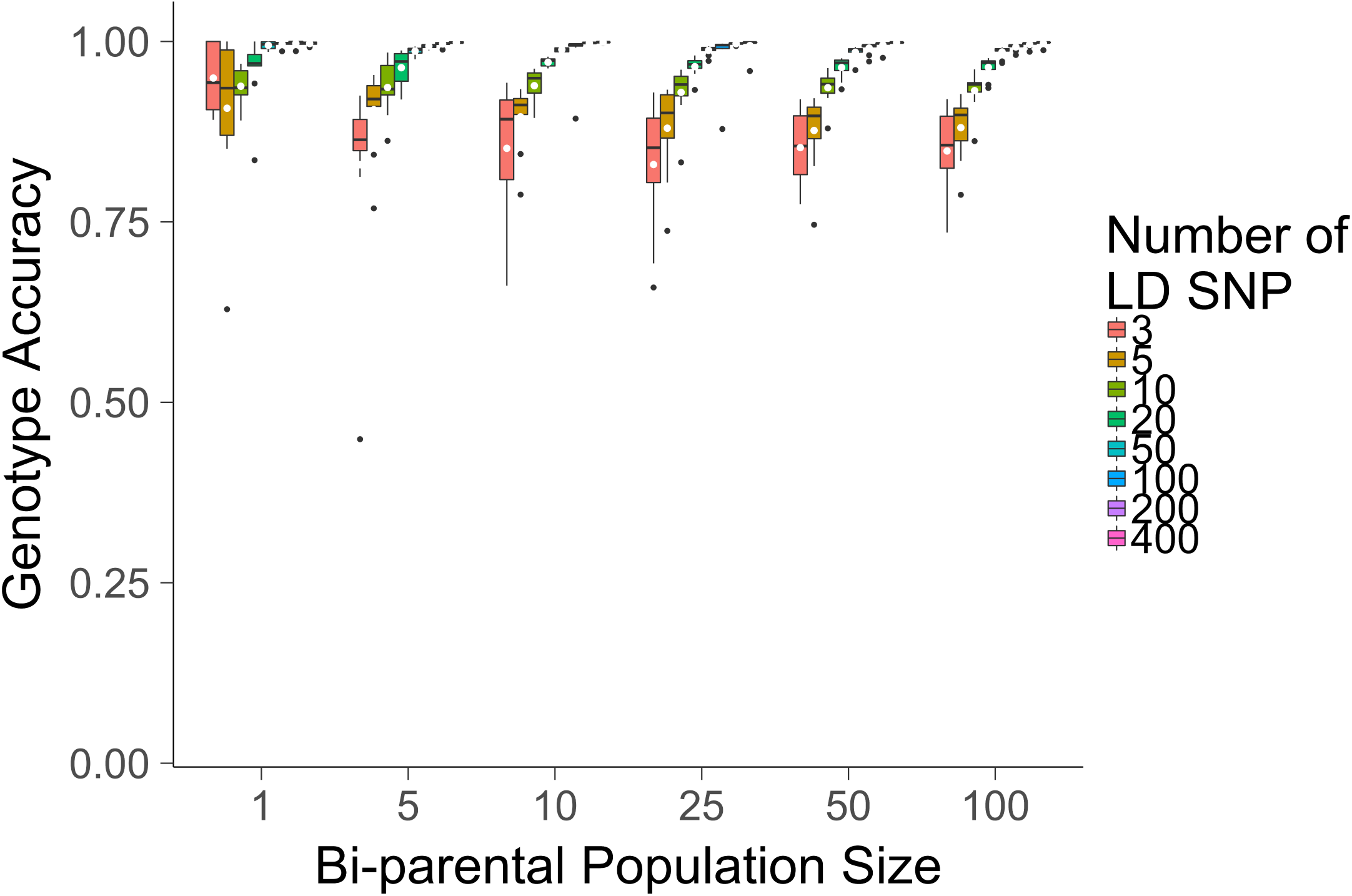
Effect of the number of focal individuals in the bi-parental population. The number of focal individuals in the bi-parental population against the genotype imputation accuracy using AlphaPlantImpute for F_2_ focal individuals of a bi-parental cross where the parents are F_20_ inbred individuals.

Figure 6 also shows that when we only imputed one focal individual, the imputation accuracy fluctuated according to the focal individual that was sampled. As a result, increasing the number of LD markers did not always increase the imputation accuracy. For example, the average imputation accuracy was 0.95, 0.91, or 0.94 when 3, 5, or 10 LD markers were used. When 400 LD markers were used, the average accuracy of imputation was 0.997.

### Computational requirements of AlphaPlantImpute

Table 1 summarises the computational requirements of AlphaPlantImpute for twelve datasets across the three scenarios. Datasets were chosen to reflect the extremes in the number of selfing events separating parents and focal individuals (F_2_ vs. F_10_), the level of inbreeding in the parents (F_1_ vs. F_20_) and the number of LD markers (3, 50, or 400). Table 1 shows that the average run time for AlphaPlantImpute was 22.13 seconds with a maximum of 49.33 seconds. The average memory requirement for AlphaPlantImpute was 0.08 GB with a maximum of 0.082 GB.

**Table 1.** Computational requirements of AlphaPlantImpute

| Parents | Focal Individuals | LD panel | Time (Seconds) | Memory (Gb) |
| --- | --- | --- | --- | --- |
| F20 | F2 | 3 | 37.41 | 0.079 |
| F20 | F2 | 50 | 8.14 | 0.080 |
| F20 | F2 | 400 | 7.95 | 0.082 |
| F20 | F10 | 3 | 49.33 | 0.079 |
| F20 | F10 | 50 | 8.48 | 0.080 |
| F20 | F10 | 400 | 9.40 | 0.082 |
| F1 | F2 | 3 | 26.70 | 0.080 |
| F1 | F2 | 50 | 35.10 | 0.080 |
| F1 | F2 | 400 | 12.58 | 0.082 |
| F1 | F10 | 3 | 24.66 | 0.079 |
| F1 | F10 | 50 | 35.41 | 0.080 |
| F1 | F10 | 400 | 10.35 | 0.082 |
| **Average:** |  |  | **22.13** | **0.080** |

## Discussion

Our results highlight three points for discussion: (i) the performance of AlphaPlantImpute; (ii) the performance of AlphaPlantImpute compared to PlantImpute; and (iii) future development of AlphaPlantImpute.

### Performance of AlphaPlantImpute

This paper presents a new heuristic method, called AlphaPlantImpute, for phasing and imputation of SNP array data in diploid plant species. AlphaPlantImpute explicitly leverages features of plant breeding programs to impute LD focal individuals to HD. The explicit utilisation of pedigree information and heuristics developed specifically to track the inheritance of parental haplotypes using the LD genotypes of focal individuals are likely to be the reasons for AlphaPlantImpute’s robust and consistent performance across all tested scenarios. AlphaPlantImpute achieves high imputation accuracy of between 0.8 and 1.0 for the majority of scenarios. For scenarios where the imputation accuracy was below 0.8, increasing the number of LD markers increased the imputation accuracy.

Increasing number of selfing events separating parents and focal individuals from F_2_ to F_10_ only slightly decreases the imputation accuracy. Decreasing the level of inbreeding in the parents or increasing the chromosome size decreases the imputation accuracy when the number of LD markers is 10 or less. However, in both cases, the decrease in the imputation accuracy could be mitigated by increasing the number of LD markers to 20 SNP or more.

Decreasing the number of focal individuals in the bi-parental population slightly decreases the imputation accuracy. This was most evident when the number of LD markers was 10 SNP or less. The likely cause of this is that inferring the most likely linkage between alleles for two markers is difficult with fewer focal individuals, since fewer individuals will be homozygous at the markers. In this case, the algorithm defaults to the linkage pattern of alleles in the parents. This may be sub-optimal for imputing markers in regions with elevated recombination rates, i.e., hotspots. When there was a single focal individual in the focal family, the accuracy of imputation for that individual varied. The likely cause of this is whether an individual had a recombination or whether it had inherited the parental haplotypes without recombination. One solution to this situation could be to utilise the most likely linkage from related families with more genotyped focal individuals (see section: Future work and developments).

Overall, the results suggest that for a given population, high imputation accuracy can be achieved even when the number of LD markers is low, and small increases in the number of markers can achieve high accuracies depending on the biology of the species (i.e. recombination rate, obligate outcrossing) and the pedigree design (outbred, inbred, level of selfing).

### Performance of AlphaPlantImpute compared to PlantImpute

The imputation accuracy for AlphaPlantImpute was compared to that for PlantImpute (Nettelblad et al., 2009; Hickey et al., 2015). In the majority of cases, the imputation accuracy was higher for AlphaPlantImpute than for PlantImpute. One exception to this was when the chromosome size was 400 cM and when the number of LD markers was 20 or less (e.g. 0.88 vs. 0.90 when the number of LD markers was 20). One reason for this could be that unless there is enough information in the genotypes of focal individual on the LD array, the heuristic algorithm in AlphaPlantImpute is inherently more conservative in determining recombination regions compared to the probabilistic algorithm in PlantImpute. As such, AlphaPlantImpute is more likely to leave positions as missing and fill them in as the parent average in the final step.

The precision of imputation accuracy (calculated as the log of the inverse of the variance in imputation accuracy within each bi-parental population) was also higher in the majority of cases for AlphaPlantImpute than for PlantImpute. This was most apparent with small number of LD markers. The higher precision of imputation accuracy for AlphaPlantImpute is likely a consequence of directly calling allele phase and parent-of-origin and imputed genotypes in turn. The probabilistic algorithm of PlantImpute is marginalizing over the all possible phase and genotype, which is probabilistically correct and handles the uncertainty properly, but it seems this is lowering the imputation accuracy. One exception to this was when the chromosome size was 150 cM, where the precision of imputation accuracy was higher for PlantImpute than for AlphaPlantImpute for all LD arrays.

The biggest advantage of AlphaPlantImpute compared to PlantImpute relates to computational requirements. Hickey et. al. 2015 report that to perform imputation within a single bi-parental population of 100 F_2_ focal individuals, PlantImpute required a minimum of 3 hours and in excess of 100 GB of memory. In comparison, AlphaPlantImpute required on average ∼22 seconds and ∼0.08 GB of memory for all tested scenarios.

The high and consistent accuracies achieved with very low computational requirements makes AlphaPlantImpute an attractive, reliable and practical tool for routine use in plant breeding programs that are already using or will include SNP array data to inform selection decisions.

### Future work and developments

At present, the heuristic method in AlphaPlantImpute works within the most common plant breeding program design of bi-parental populations and it works best when parents are fully inbred or close to being fully inbred. AlphaPlantImpute could be extended in multiple ways. For example, instead of treating each bi-parental population as an independent unit it could simultaneously work across bi-parental populations that share parents. This could increase the imputation accuracy in three ways: (i) information between bi-parental populations could be shared for imputation of focal individuals that are effectively half-sibs (one common parent); (ii) information between bi-parental populations could be used to resolve phase where one or both parents are heterozygous at one or more consecutive markers; and (iii) if a common parent has no or LD genotypes available, information from its descendants across half-sib bi-parental populations could be leveraged to phase and impute it to high-density.

AlphaPlantImpute could also be extended to include ancestral pedigree information (such as grandparents and great-grandparents). This could be useful for improving phasing and imputation of parents with missing information or that are highly outbred. More simply, AlphaPlantImpute could also be extended so that it can directly read in and exploit phased information for the fully or partially outbred parents. Such phased information could be generated for parents by running AlphaPlantImpute on the bi-parental family from which the fully or partially outbred parent derived.

AlphaPlantImpute could be extended so that it reads in previously inferred most likely linked alleles at two markers. It is likely that linkage patterns are shared across families, especially if the families are related. Using this information across families would be especially suited to imputation situations in bi-parental populations that have only a few genotyped focal individuals (e.g., one genotyped individual per family).

Finally, although SNP arrays for the many domesticated plant species exist, low-coverage sequencing methods such as genotyping-by-sequencing are also used. The heuristics of AlphaPlantImpute might be extended to enable imputation with such data.

### Software availability

We implemented our method in a software package called AlphaPlantImpute, which is available for download at http://www.AlphaGenes.roslin.ed.ac.uk/AlphaPlantImpute/ along with a user manual.

## Acknowledgments

The authors acknowledge the financial support from the BBSRC ISP grant number ‘BB/P013759/1’ and from the BBSRC KWS grant number ‘BB/R002061/1’. This work has made use of the resources provided by the Edinburgh Compute and Data Facility (ECDF) (http://www.ecdf.ed.ac.uk).

## Supplementary Files

**Supplementary File 1. Detailed schematic of heuristic algorithm of AlphaPlantImpute.**

## Abbreviations

LD: low-density
HD: high-density
SNP: single nucleotide polymorphism
cM: centiMorgan

